# Derivation of oligodendrocyte precursor cells from human bone marrow stromal cells and use for re-myelination in the congenitally dysmyelinated brain

**DOI:** 10.1101/658997

**Authors:** Guy Lam, Graham Ka-Hon Shea, Lap Kei Wu, Maximillian Li Tak Sui, Kwok Chun Hei, Zora Chan Chui Kuen, Yvonne Wong Cheuk Yin, Alex Yat Ping Tsui, Daisy Kwok Yan Shum, Ying Shing Chan

## Abstract

Oligodendrocytes (OLs) are the only myelinating glia in the central nervous system (CNS). In congenital myelin disorders, OL dysfunction or death results in loss of myelin. This causes progressive and irreversible impairment to motor and cognitive functions, and is amongst the most disabling neurological disorder.

Neonatal engraftment by glial progenitor cells (GPCs) allows the robust myelination of congenitally dysmyelinated brain, thereby preserving brain function and quality of life of patients. However, endogenous sources of glial progenitors are hard to obtain without causing secondary injury, while use of exogenous sources such as embryonic stem cells and induced-pluripotent stem cells face considerable ethical and safety issues.

To circumvent such hurdles, we asked whether NG2^+^ cells in the bone marrow could be a potential cell source for GPCs. We successfully generated glial progenitor cells (GPCs) from human bone marrow stromal cells (hBMSCs) from 3 donors using a 14- day induction protocol. The generated hBMSC-GPCs were highly enriched in OPC marker expression, including OLIG2, PDGFRα, NG2, SOX10 and O4, and showed efficient differentiation into myelinogenic oligodendrocytes when transplanted into postnatal day 7 (P7) myelin-deficient shiverer mice. Remyelination of the shiverer mouse brain significantly extended lifespan and improved motor function.

The novel induction protocol described here provides a method for fast, simple and effective glial therapy for myelin disorders, overcoming existent hurdles of cell source restriction and time frame requirement.

**Graphical Abstract:** 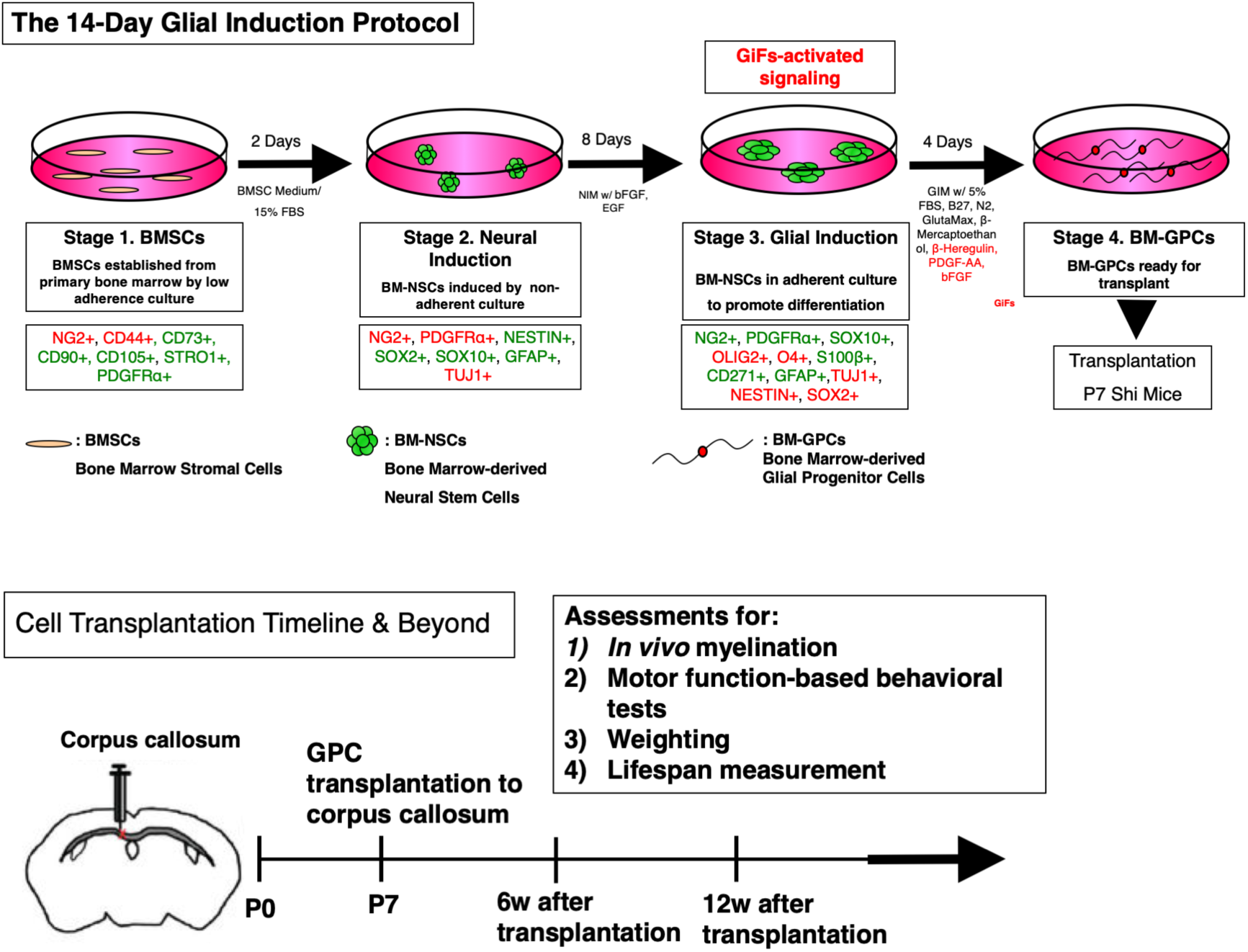

**Highlights:**

- Human bone marrow stromal cells (BMSCs) can be programmed to myelinating glia (GPCs, glial progenitor cells), via a novel 14-day *in vitro* induction protocol
- Transplantation of these hBM-GPCs robustly remyelinated myelin deficient shiverer mice.
- hBM-GPC transplant significantly extended lifespan, increased body weight and improved motor function

## INTRODUCTION

### The significance of myelination

Ensheathment of axons by myelin in the white matter provides electrical insulation for the fast and effective transmission of nerve impulses, as well as protection for axons against inflammatory and oxidative insults.^1^ (Bankston et al., 2013)

In the Central Nervous System (CNS), oligodendrocytes (OL) are responsible for myelinating axons.^2^ (Baumann et al., 2001) Myelin disorders due to OL dysfunction or death, resulting in a loss of myelin. Myelin disorders are amongst the most disabling and prevalent of neurological conditions. Myelin disorders can be classified into (1) myelinoclastic diseases and (2) demyelinating leukodystrophic diseases ^3^ (Love, 2016) and (3) demyelination due to insult, such as radiation therapy to the brain or exposure to OL-toxins such as ethidium bromide. Myelin disorders cause a progressive and irreversible impairment in cognitive function as well as motor and executive impairment due to the depletion of the oligodendrocyte precursor pool and subsequent demyelination. ^4^ (Piao et al., 2015)

### Demyelination, cell replacement therapy and application

Rapid remyelination of demyelinated axons is crucial for preservation of axon connectivity and the function of neural circuitries, since demyelinated axons not only have lower conductance, they are vulnerable to degenerative processes.

The ability to harness and commit stem cells to specific neural fates with a high reproducibility has opened the doorway to effective cell replacement therapies. (Tabar and Studer, 2014)^7^ Various cell-based strategies have been established for repair of demyelinated lesions in both the brain and spinal cord (Ben-Hur and Goldman, 2008; Franklin and ffrench-Constant, 2008).^5, 6^

Congenital genetic disorders such as myelin basic protein defect-associated dysmyelination, where endogenous glial cells and their progenitors are not available/fail to perform effective myelination (Franklin and ffrench-Constant, 2008) ^6^, are compelling targets for cell replacement strategies. Aside from overt OL loss and failure to form myelin (such as in leukodystrophies and inflammatory demyelinations), several neurodegenerative and psychiatric disorders may be linked to astrocytic and oligodendrocytic pathology causally as well. (Goldman et al., 2017) ^8^ The relative contribution of glial dysfunction to these disorders may allow corresponding treatment by the transplantation of allogeneic glial progenitor cells (GPCs), the precursors to both astroglia and myelin-producing OLs.

### Glial Progenitor Cells and replacement strategies

#### I) *In Vivo* Role of GPCs

Human neuron-glial antigen 2 (NG2)-positive glial progenitor cells (GPCs) arising from neural stem cells of the ventricular subependyma. (Nishiyama et al., 2009; Goldman et al., 2012) ^9,10^, are found throughout the brain in both gray and white matter (Roy et al., 1999) ^11^. *In vitro* and *in vivo* studies have demonstrated that human GPCs are bipotent in nature until the last division, generating both major macroglial phenotypes, astrocytes and OLs, in a context-dependent fashion (Windrem et al., 2004)^12^. As the principal source of new OLs, GPCs are responsible for translational remyelination in the demyelinated adult CNS (Tripathi et al., 2010; Zawadzka et al., 2010) ^13,14^.

#### II) Translational application

As such, GPCs can concurrently restore myelin, while addressing the astrocytic function disorders related to white matter disease. (Goldman et al., 2017) ^8^

Since harvesting primary GPCs from neural tissue induces secondary damage, recent studies on GPC transplant employed human (1) embryonic stem cells, (2) fetal and adult brain tissues and (3) induced pluripotent stem cells (iPSCs) as the source of GPCs. (Dietrich et al., 2002; Roy et al., 1999; Windrem et al., 2004; Hu et al., 2009; Izrael et al., 2007; Keirstead et al., 2005) ^15,11,12,16,17,18^. GPCs thus derived were shown to mitigate both congenital (Sim et al., 2011; Windrem et al., 2004; Windrem et al., 2008; Duncan et al., 2009; Hu et al., 2009, Wang et al., 2013; Douvaras et al. 2014; Yang et al. 2013) ^19, 12, 20,16, 21, 22, 23^ and adult (Windrem et al., 2002) ^24^ CNS demyelination. However, use of ESCs and iPSCs face significant ethical and safety concerns (Sugarman et al., 2008; Saha et al., 2009) ^26, 27^.

Additionally, while existing GPC derivation protocols have been proven to yield functional myelinating glia, they remain complex and protracted. To the best of our knowledge, the fastest protocol requires at least 28 days and viral media to generate GPCs (Ehrlich et al., 2017) ^28^, while others usually require months. (Piao et al., 2015; Wong et al., 2013) ^4, 21^ This limits therapeutic usage for adult demyelination. It highlights the importance and benefit of developing a fast and robust method for obtaining GPCs from a safe and readily available cell source to meet the demands of clinical application.

### Our approach on cell replacement therapy

Adult bone marrow stromal cells (BMSCs) represent a potential, safe and ready source of neural progenitors for autologous transplantation devoid of the previous concerns (Tomita et al., 2006; Chen et al., 2006) ^29, 30^. In this regard, we aimed to convert BMSCs into GPCs with a faster, non-viral protocol of higher yield.

### PDGF signaling & Glial fate commitment

Since the last three decades, Platelet-derived growth factor (PDGF) signaling has been known to positively regulate: 1) migration, 2) proliferation, and 3) differentiation of glial progenitors during brain development. (Noble et al., 1988; Armstrong et al., 1990; Frost, Zhou, Krasnesky, & Armstrong, 2009; Assanah et al., 2009) ^47, 48, 52^

PDGF has been previously shown to drive massive expansion of glial progenitor cell in the Neonatal Forebrain (Assanah et al., 2009) ^54^. PDGF-AA (alpha form) molecules not only serve as a chemoattractant for glial progenitor migration (Armstrong et al., 1990)^48^, but when combined with fibroblast growth factor (FGF) in culture, these progenitors cells remain immature, migratory, and proliferative for an extended period of time (Bogler et al., 1990; Tang et al., 2001). ^49, 53^

In terms of cell signalling, the mitogenic activities promoted by combined PDGF/ FGF-2 signaling is mediated through activation of the MAPK/ERK pathway, specifically involving: p42/p44 mitogen-activated protein kinase (MAPK), p38 MAPK, and pp70 S6 kinase. (Baron, Metz, Bansal, Hoekstra, & De Vries, 2000)^51^. This also hinted that molecular mechanism controlling oligodendrocyte lineage progression is directed by at least two signal pathways, which either regulates proliferation and/or differentiation of oligodendrocyte progenitors. (Bogler et al., 1990; Tang et al., 2001) ^49, 53^

In a translational perspective, this discovery is particularly favourably since: 1) mature oligodendrocytes exhibit arrest and death in the absence of axonal support in culture & 2) the migratory properties is exceptionally considerable to enhance widespread remyelination upon transplantation.

### Our Findings

Human BMSCs from 3 traumatically injured patients were used generate glial progenitor cells (GPCs) in a 14-day GPC induction protocol which (i) fostered expansion of neuroprogenitor cells in sphere-forming suspension culture and (ii) following transfer to adherent culture, directed differentiation of the neuroprogenitor cells along the central glial lineage with use of a cocktail of beta-heregulin, PDGF-AA, and bFGF. A high percentage of the generated cell population express OPC markers (OLIG2, PDGFRα, NG2, SOX10 and O4), supporting the potential of BMSCs to be programmed into myelinating glia. Neonatal engraftment of GPCs derived using our protocol into myelin-deficient Shiverer mice resulted in remyelination of axons in the brain. GPC transplant ameliorated sensory-motor deficits and extended the lifespan of these mice.

This study thus provides a novel induction protocol and a new viable human cell source for prospectively autologous, fast, simple and clinically safe glial therapy to treat both congenital and acquired myelin disorder, overcoming existing hurdles of cell source restriction and time frame requirement.

## RESULTS

### Human BMSCs efficiently programmed to glial cell fate in 14 days

The first question we asked, with reference to current limitations of OPC therapies, is whether mesenchymal stem cells (MSCs) could be directed into glial fate within a short timeframe for clinical application.

To this end, we developed a 3-stage protocol to generate early myelinating glial progenitor cells within 14 days (detailed in Supplemental Methods and schematized in Figure 1**A**), comparing to past protocols that require 110-150 days (Wang et al., 2012)^21^.

**Figure 1.**
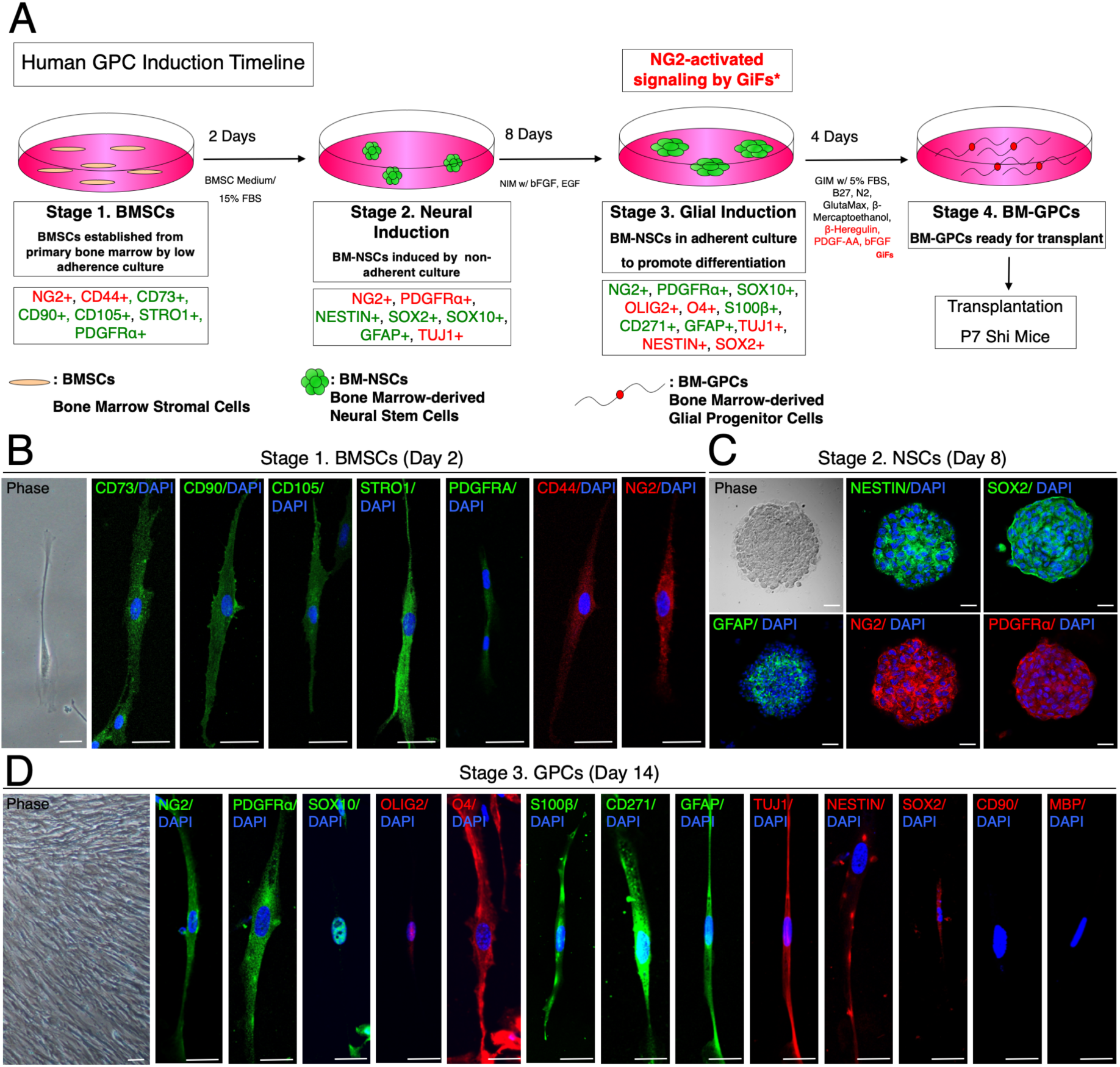
Direction of hBMSCs into GPC fate within 14 days. (**A**) A schematic protocol for directed differentiation of hBMSCs into GPCs in 14 days. BMSCs from human patients (hBMSCs) were purified and established *in vitro* from Day 0-2. Spheres of neural stem cells (NS; neurospheres) were differentiated from hBMSCs (Stage 1) from Day 3-10 in neural induction medium (NIM; refer to Methods) with bFGF and EGF. From Day 11-14, neurospheres were then pushed towards glial fate induction to become GPCs in glial induction medium (GIM; refer to Methods) featuring critical Glial induction Factors (GiFs: β-Heregulin, PDGF-AA, bFGF). (**B**) Immunocytochemistry of Stage 1, undifferentiated hBMSCs. MSC markers CD90+/ CD105+/ CD44+/CD73+/ STRO-1+ were expressed. NG2, PDGFR*α* were also expressed. (**C**) By Stage 2, the spheres expressed neural stem cell markers, being NESTIN+/ SOX2+/ GFAP+/STRO-1+. NG2, PDGFR*α* was also expressed. (**D**) By Stage 3, markers of early myelinogenic glial progenitors, including: NG2/ PDGFRα/ OLIG2/ SOX10/ O4 were highly expressed. Neural stem cell markers: SOX2/ NESTIN/ TUJ1 were also present. Astrocyte marker GFAP and PNS glial markers: CD271/ S100β were also expressed. MSC marker CD90 & mature OL marker: MBP were not expressed, indicating the early myelinogenic fate of the derived cell. Scale: **B, C, D**: 25 μm.

Human bone marrow stromal cells (hBMSCs) harvested from 3 donors were used as the starting source for GPC generation. All three bone marrow samples were harvested from the tibia of patients that suffered from bone fracture across a great age variety (Age: 11, 59, 82 Years) with no other disease diagnosed.

A population of mesenchymal stem cells (MSCs), as indicated by immunoreactivity for CD90^+^/ 105^+^/ 44^+^/ 73^+^/ STRO-1^+^/ NG2^+^/ PDGFR*α*^+^, was first isolated from the sample by selective adherent culture (Figure 1**B**). These cells were then harvested and expanded in suspension induction culture for neurospheres. After 8 days of neural induction, intermediate neural stem cells from neurospheres were highly enriched in neural stem cell markers NESTIN, SOX2 (Figure 1**C**) and GFAP.

8-day old neurospheres were then returned to adherent culture for glial induction. At the 4^th^ day of the glial induction, NG2^+^/ PDGFRα^+^/ SOX10^+^/ OLIG2^+^/ O4^+^ was highly expressed in the cell population, with suppressed NESTIN and SOX2 (neural stem cell markers) expression, across all three patients.

The generated glial progenitor cells (GPCs) possess a tripotential fate to be differentiated into astrocytes, oligodendrocytes (OLs) or neurons, shown by the simultaneous expression of both astrocyte, OPC and neural stem cell markers within the same population (Figure 1**D**, Supplementary Figure 1).

### Derived hBM-GPC population highly expressed OPC markers

To quantify myelinogenic glial progenitor purity within our obtained GPC culture, flow cytometry analysis was conducted to characterize the derived cell population.

Undifferentiated hBMSCs of the human samples consistently expressed MSC markers to a high extent (>90%) (Figure 2**A**), as follows: CD90^+^: 99.43%, CD105^+^: 97.73%, CD44^+^: 88.36%, CD73^+^: 98.23%, STRO-1: 96.4% and Alpha SMA: 99.45%. NG2 & PDGFR*α* was also expressed up to 89.8% & 73.53% (n= 3, 16,000-19,000 gated events per sample). This is in line with immnostaining data reported in Figure 1**B**.

**Figure 2.**
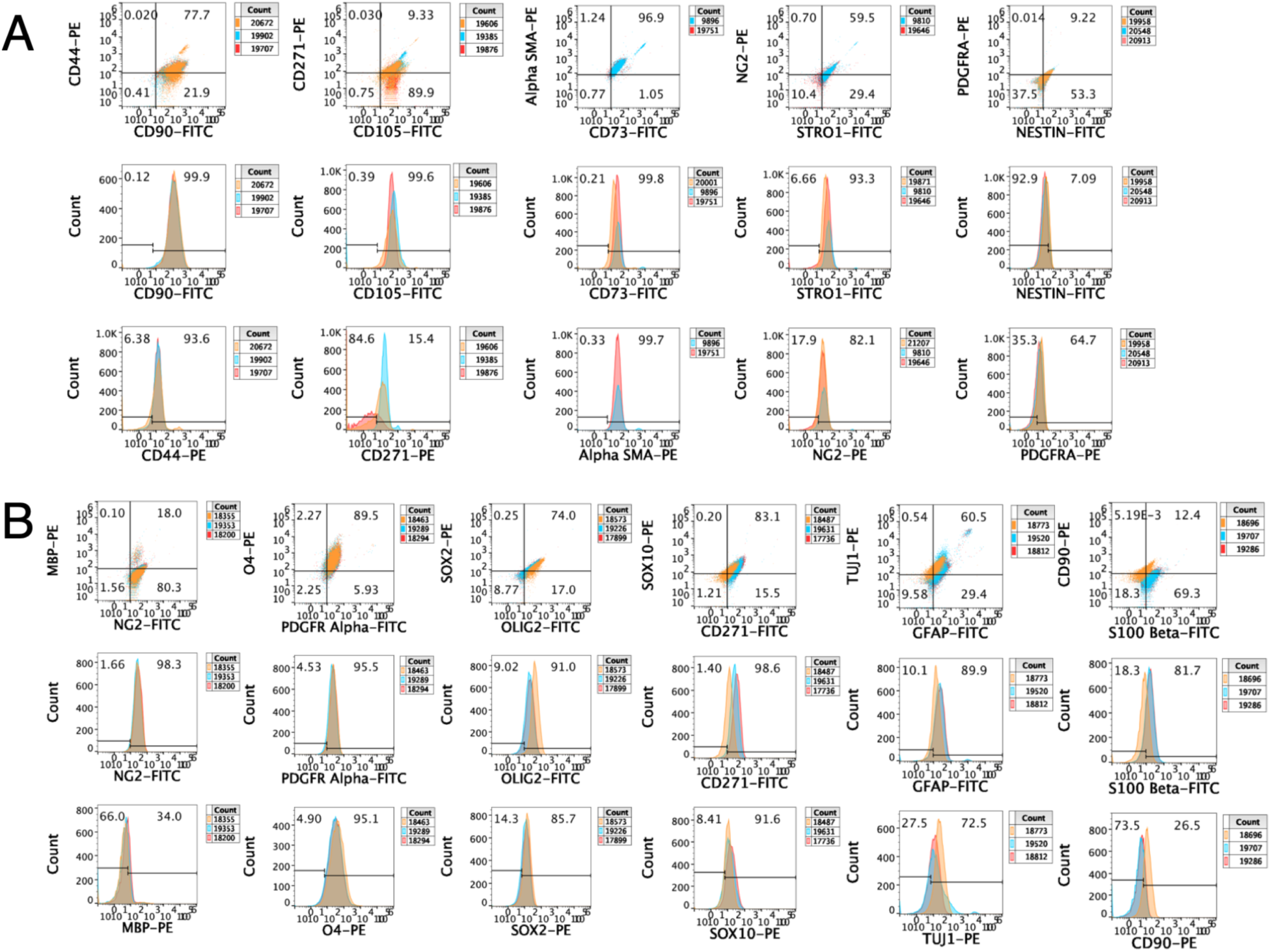
Induced GPCs feature high myelinogenic glial maturation potential. (**A**) Undifferentiated hBMSC population (n=3, labelled respectively in Red, Blue & Orange, with ∼20,000 gated events per sample) was characterized with flow cytometry. BMSC markers CD44, CD73, CD90, CD105, STRO-1, Alpha SMA were consistently expressed in more than 90% of cells; NG2 & PDGFR*α* was also expressed up to 82.1% & 64.7%. (**B**) Generated hBM-GPC population (n=3, labelled respectively in Red, Blue & Orange, with ∼20,000 gated events per sample) was characterized with flow cytometry analysis. Over 90% of the population expressed OPC markers NG2, PDGFRα, O4 and OLIG2 in concordance with immunocytochemistry data presented in Figure 1, suggesting their potency to generate OLs. Neural stem cell, astrocyte and PNS glial potency was also suggested by the expression of TUJ1, GFAP, CD271 and S100β.

Generated hBM-GPC population highly expressed OPC markers: 98.27% NG2+, 95.77% PDGFRα+, 94.13% O4+, 93.16% OLIG2+, 75.8% SOX10+ (Figure 2**B**, n=3, labelled respectively in Red, Blue & Orange, with ∼20,000 gated events per sample). This also corroborated immunostaining data and suggested potency of our derived cells in generating OLs. The majority of the generated GPCs remained as progenitors as shown by the low expression of the mature OL marker myelin basic protein (MBP). Notably, the cells retained neural stem cell (TUJ1: 78.23%), astrocyte (GFAP: 89.5%) and PNS glial (S100β: 74.23%, CD271: 94.1%) potency. With respect to the overall high purity of cells by marker selection, no specific cell sorting was required.

*(Whole Transcriptome Sequencing to be supplemented here upon completion)*

### Neonatally transplanted hBM-GPCs functionally myelinated the brain *in vivo*

Given the high purity of GPCs generated, as suggested by marker expression, we asked if the cells could indeed generate mature myelinating OLs in vivo. To address this question, we transplanted the cells into newborn homozygous shiverer mice at postnatal day 7 (P7), taking advantage of immune tolerance in the early postnatal stage. Neonatal pups were transplanted bilaterally in the corpus callosum (1mm next to the central cleavage of the corpus callosum, 1.5mm deep into cortex) and the hindbrain with a total of 90,000 cells, with reference to previous studies (Windrem et al., 2008; Wang et al., 2013) ^20, 21^ At week 12, the mice were sacrificed for analysis of donor cell localization, density, myelination patterns and myelination efficiency.

hBM-GPCs generated from all 3 donors were able to myelinate the shiver brain 11 weeks post-transplant, as shown by confocal images of week 12 shiverer mice brain coronal sections stained for MBP (Figure 3**B**).

**Figure 3.**
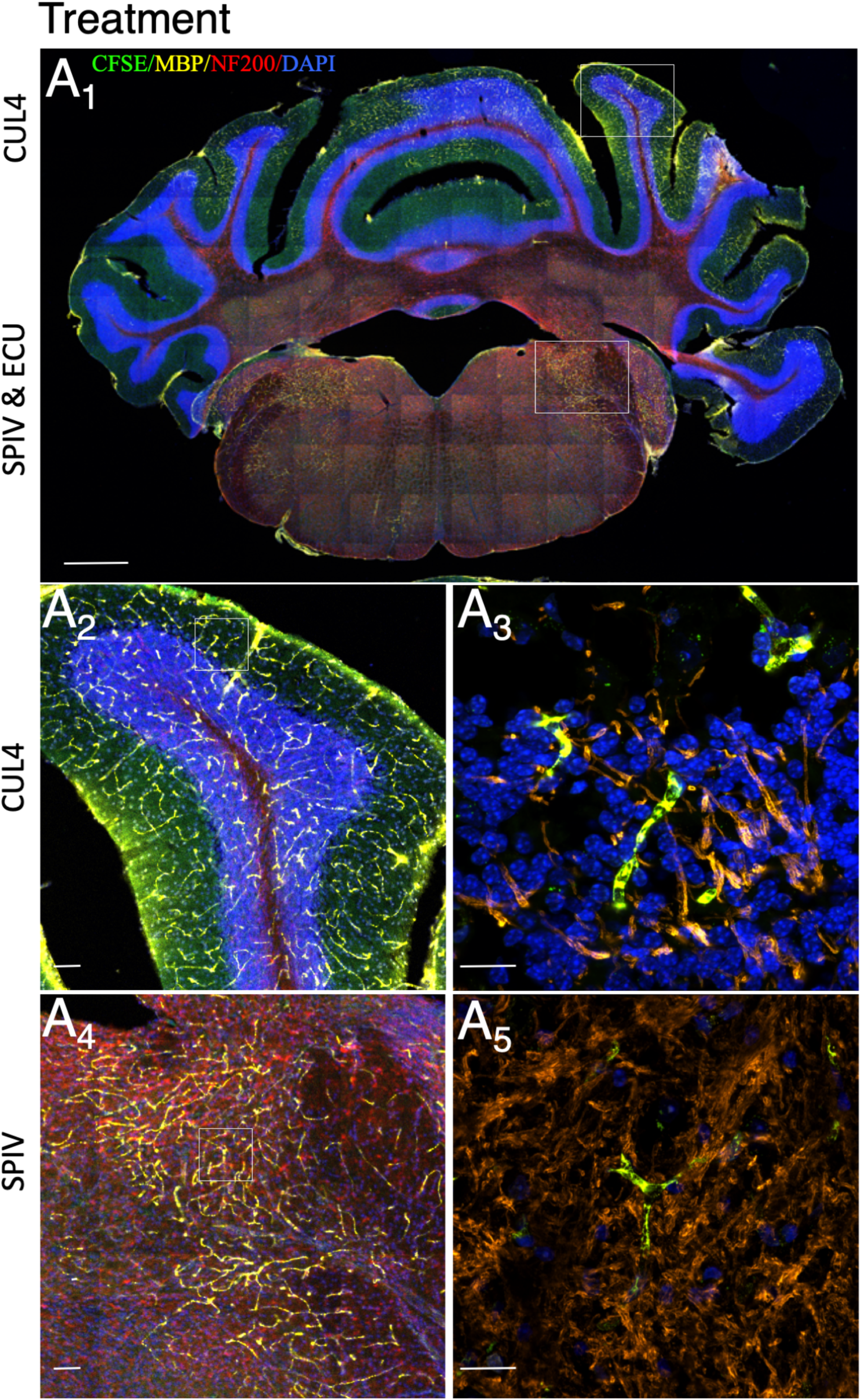
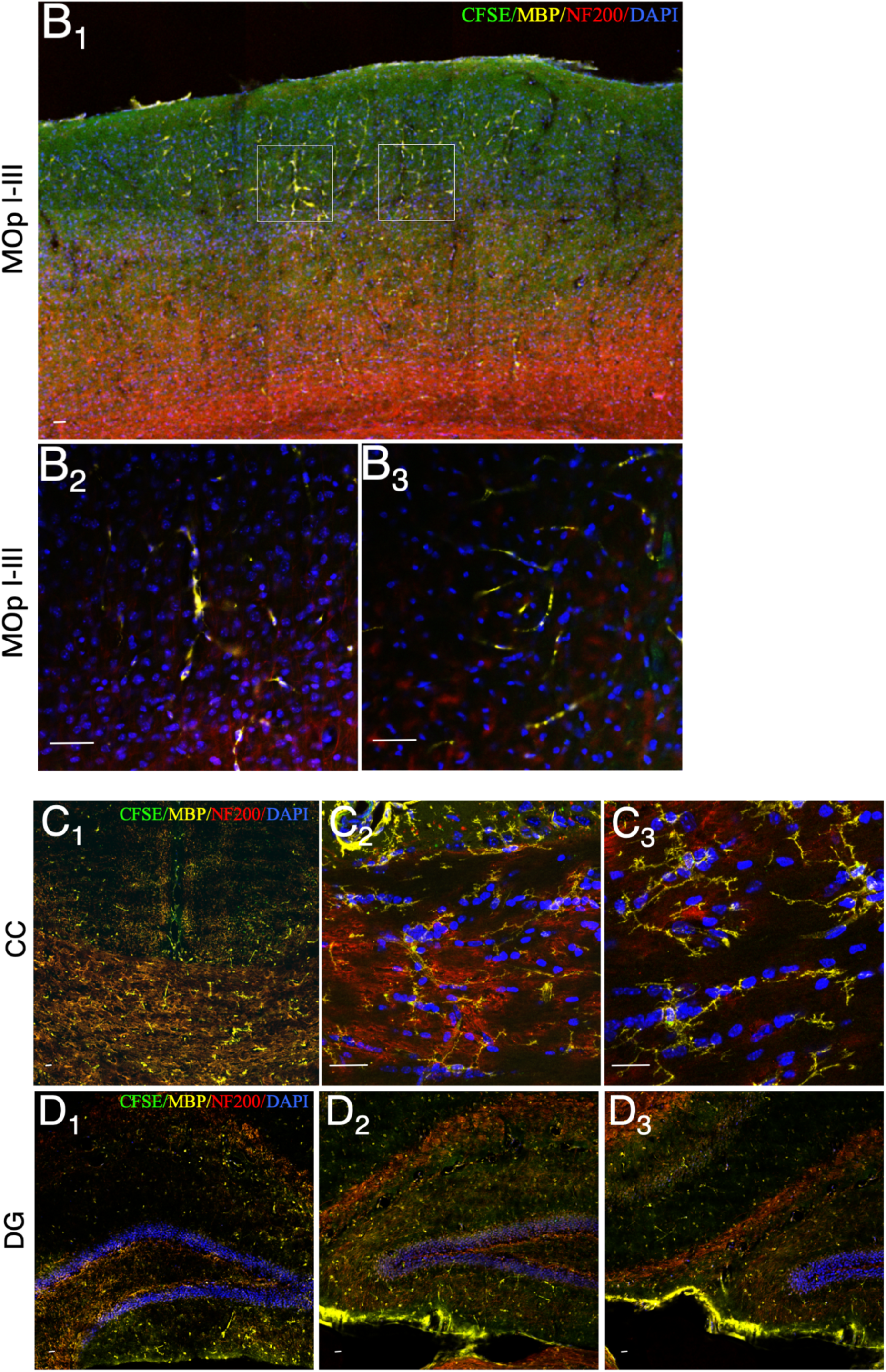
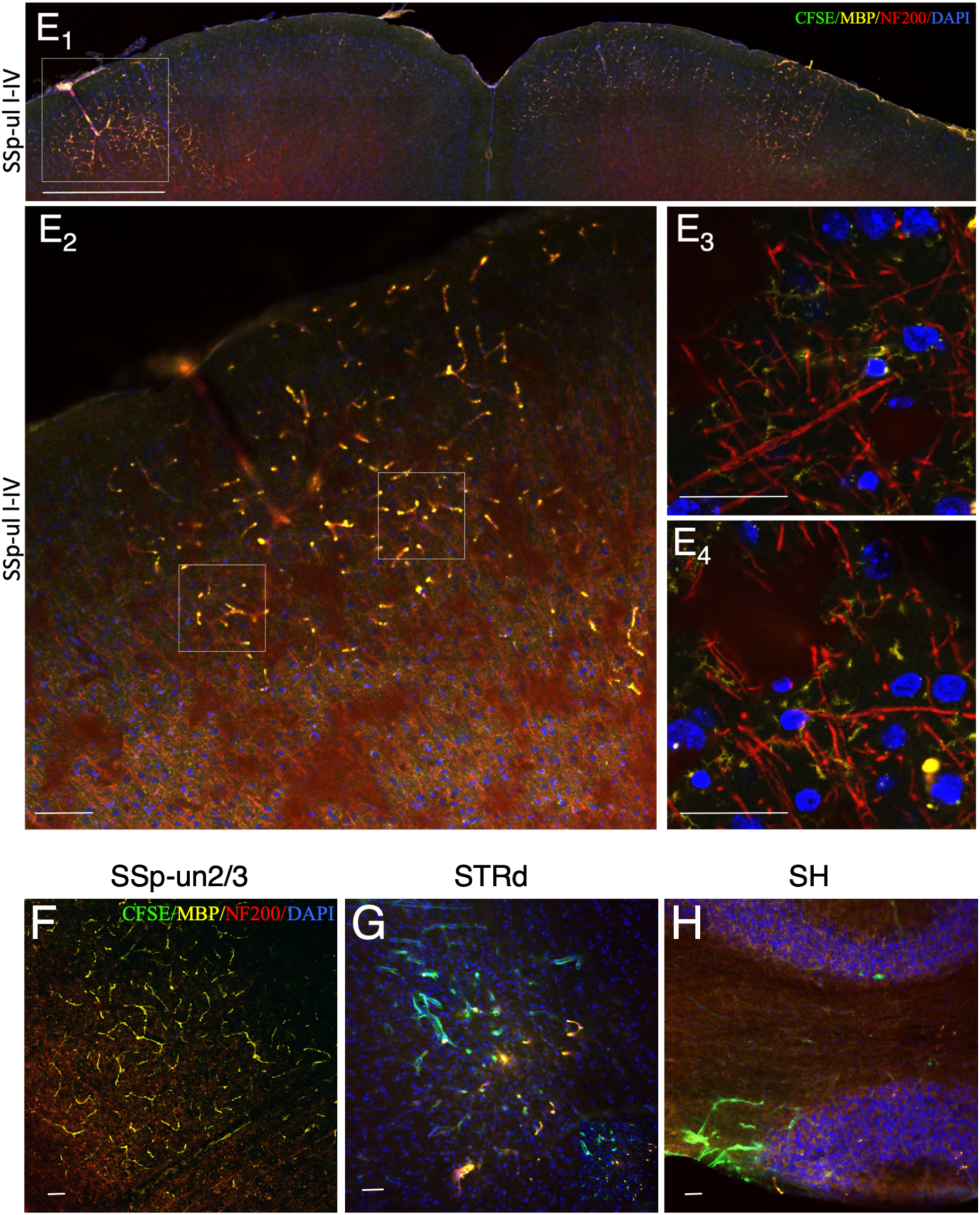
IHC Confocal Images of hBM-GPCs-derived oligodendrocytes ensheathed axons in shiverer brain. Confocal images of coronal sections of the shiverer mice brain harvested at week 12. (**A**_**1**_) Histology featuring the coronal hind-brain section of a treated shiverer mouse. Widely dispersed myelination is featured, extensive myelination locates at the spinal vestibular nucleus (SPIV), external cuneate nucleus (ECU) and cerebellar lobules (CUL4 IV-V). (CFSE of human cell, green; MBP, yellow, NF200, red). (**A**_**2**_, **A**_**3**_) Local Myelination (CFSE of human cell, green; MBP, yellow, NF200, red) at the cerebellum lobules (CUL4) and high power magnification (63x) showing the myelinated fine axons. (**A**_**4**_, **A**_**5**_) Local myelination (CFSE of human cell, green; MBP, yellow, NF200, red) at the spinal vestibular nucleus (SPIV) and high power magnification (63x) showing the myelinated fine axons. Confocal images of coronal sections of the shiverer mice brain harvested at week 12. (**B**_**1**_) Histology featuring the coronal hind-brain section of a treated shiverer mouse. Myelination locates at the motor cortex (MOpI-III). (CFSE of human cell, green; MBP, yellow, NF200, red). (**B**_**2**_, **B**_**3**_) High power magnification (63x) showing the myelinated axons of the motor cortex (MOpI-III). (**C**_**1**, **2**, **3**_) Extensive myelination locating at the corpus callosum (CC). (CFSE of human cell, green; MBP, yellow, NF200, red). (**D**_**1**, **2**, **3**_) Local Myelination (CFSE of human cell, green; MBP, yellow, NF200, red) at the dentate gyrus. Confocal images of coronal sections of the shiverer mice brain harvested at week 12. (**E**_**1**_) Histology featuring the coronal mid-brain section of a treated shiverer mouse. Myelination locates at the somatosensory cortex (SSp-ul I-IV). (CFSE of human cell, green; MBP, yellow, NF200, red). (**E**_**2**, **3**, **4**_) High power magnification (63x) showing the myelinated axons of the somatosensory cortex (SSp-uI-IV). Local Myelination (CFSE of human cell, green; MBP, yellow, NF200, red) at the Somatosensory Cortex (SSP-un2/3, Layer 2-3). Local Myelination (CFSE of human cell, green; MBP, yellow, NF200, red) at the dorsal Striatum (STRd). Local Myelination (CFSE of human cell, green; MBP, yellow, NF200, red) at the septohippocampal nucleus (SH). Scale: **A**_**1**_, **E1**: 1 mm; **A**_**2**_, **A**_**3**_, **A**_**4**_, **A**_**5**,_ **B**_**1**_, **C**_**1**, **2**, **3**_, **D**_**1**, **2**, **3**_, **E**_**2**, **3**, **4**_, **F, G, H**: 100 μm. Anatomical Abbreviations: CC Corpus Callosum CUL Cerebellar lobules DG Dentate Gyrus ECU External cuneate nucleus MOp Motor Cortex SH Septohippocampal nucleus SSp-un Somatosensory cortex (unassigned) SP Spinal vestibular nucleus STRd Striatum (dorsal region)

Our hBM-GPCs demonstrated extensive post-transplantation ventral (inferior) migration, along the injection site down the corpus callosum and the hindbrain.

At the posterior injection site in the hindbrain, the cells were found well-dispersed, which suggests migration throughout most of the cerebellum (Figure **3 A**_**1**_). Extensive myelination was located at the spinal vestibular nucleus (SPIV) (Figure **3 A**_**4**,**5**_), external cuneate nucleus (ECU), as well as the cerebellar lobules (CUL4 IV-V, molecular and granular layer) (Figure **3 A**_**2-3**_).

Dorsal (superior) migration was also found as myelin, along with labelled cells, at the outer layers (Layer I-III: molecular, external granular & external pyramidal layer) of the motor cortex (MOpI, II, III) (Figure **3 B**_**1-3**_) and somatosensory cortex (SSp-II1, 2/3) (Figure **3 E**).

At the two anterior injection sites, extensive ventral migration was demonstrated along the corpus callosum (CC) (Figure **3 C**_**1-3**_), as well as the dentate gyrus (Figure **3 D**_**1-3**_). Migration was found all the way to the striatum (STRd) (Figure **3 G**) and septohippocampal nucleus (SH) (Figure **3 H**), reaching as far as the lateral ventricle. In contrast, the controls (Sham and medium-injected) yielded none of the above observations (Supplementary Figure 4).

Comparatively, limited rostral-caudal migration along the striatal tract and the hindbrain was also observed in IHC sections. (+ **Elaborate**, *Number to be supplemented, quantitatively (density/ distribution) data of CFSE*^*+*^*/ MBP*^*+*^ *cells along NF200*^*+*^ *axons currently unavailable*.) Early quantitative analysis of 3D renderings (Imaris) revealed a mean density of 638 cells/1mm^2^ at the corpus callosum per 20 μm thick of section.

### hBM-GPCs-derived oligodendrocytes ensheathed axons *in vivo* in shiverer brain

To ascertain that the observed MBP signal was resultant from formation of compact myelin, ultrathin sections of corpus callosum and upper striatum tissue were obtained from 12-week old shiverer mice for electron microscopy.

Compact myelin around axons were abundant in corpus callosum tissue from hBM-GPC–transplanted mice (**Figure** 4**B**), which were absent in untreated shiverer controls (**Figure** 4**A**_**1-2**_).

The upper striatum featured rings of immaturely myelinated axons (Figure 4 **B**_**1-5**_) and at the callosa, numerous compact myelin sheaths ensheathing central axons at the corpus callosum were observed (Figure 4**B**_**6-9**_). The tissue was myelinated at a density up to 56 axons/ 500um^2^.

**Figure 4.**
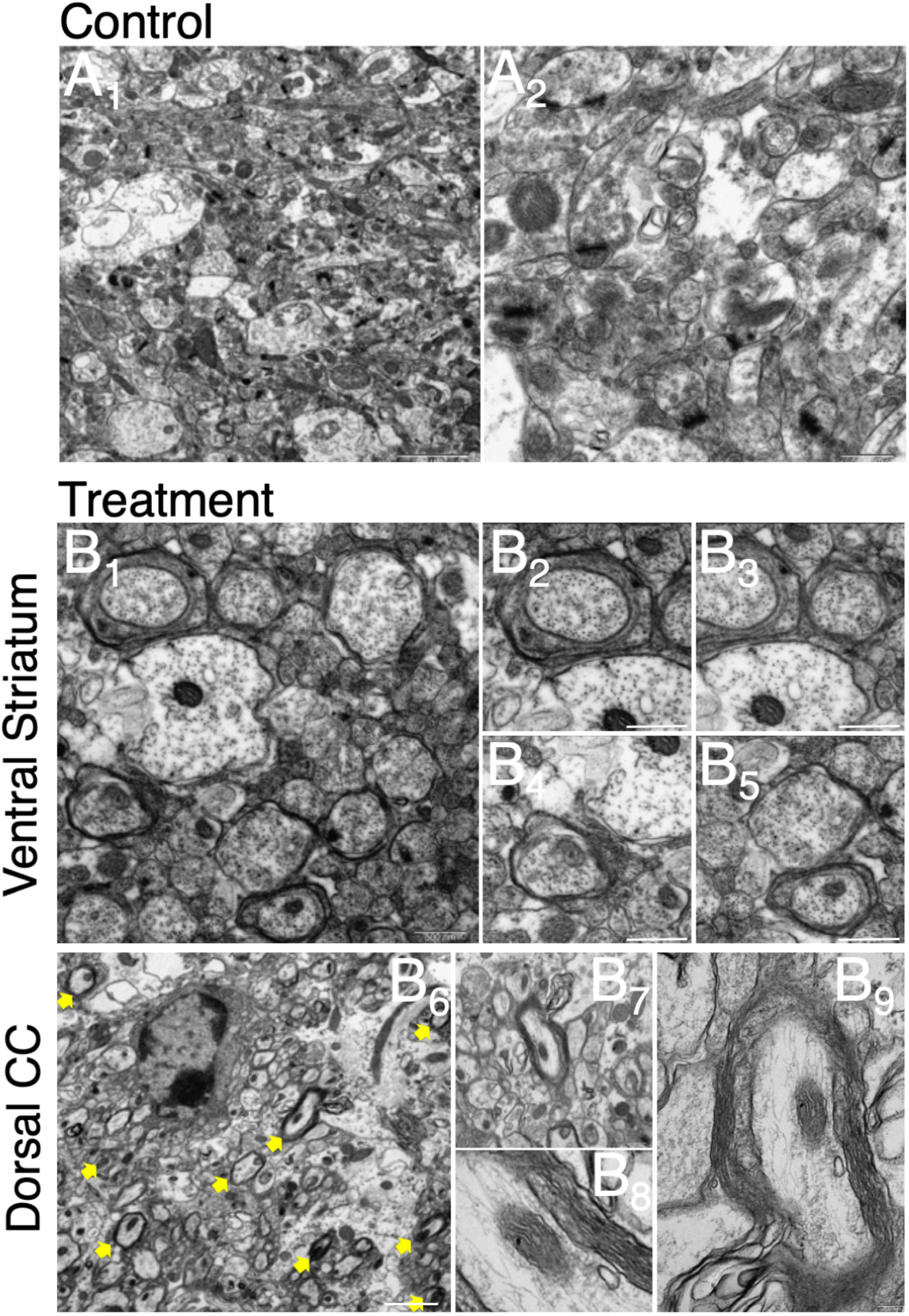
hBM-GPCs ensheathed axons in myelin ambient to the transplantation site. Representative electron microscopy images of sagittal sections of the upper shiverer mice brain harvested at week 12. (**A**) Histology featuring the unmyelinated axons of a control shiverer mice. (**B**_**1-5**_) Histology featuring a ring of immaturely myelinated axons at the upper striatum. (**B**_**6-9**_) Histology featuring multiple compact myelin sheaths ensheathing central axons at the corpus callosum. Scale: **A**_**1-2**_: 500 nm; **B**_**1**_:500nm, **B**_**2-5**_: 200 nm, **B**_**6**_: 2 um, **B**_**7**_: 500 nm, **B**_**8-9**_: 200 nm.

As a summary, hBM-GPCs-derived oligodendrocytes myelinate axons compactly *in vivo* in shiverer brain.

### Neonatally transplanted hBM-GPCs rescue motor function, body weight and extend shiverer lifespan through remyelination

We finally asked if myelination of corpus callosum and striatal tissue could improve symptoms resultant from congenial loss of myelin. Body weight and motor function were assessed in all the mice at week 6 and week 12. The lifespan of mice with and without hBM-GPC transplantation was also recorded. (Detailed in Supplemental Methods)

Cell-transplanted (treatment group) animals were significantly heavier than controls at 12 weeks. a timepoint featuring completion of murine maturational growth, shiverer mouse functional decline, arrest and death. (Figure 5**A**_**2**_) A 13.98% increase (27.48g to 20.89g) for the week 12 male group and 14.5% increase (23.92g to 20.89g) for the female group was observed. At week 6, a timepoint corresponding to adolescence, no significant difference in body weight was found. (Figure 5**A**_**1**_)

**Figure 5.**
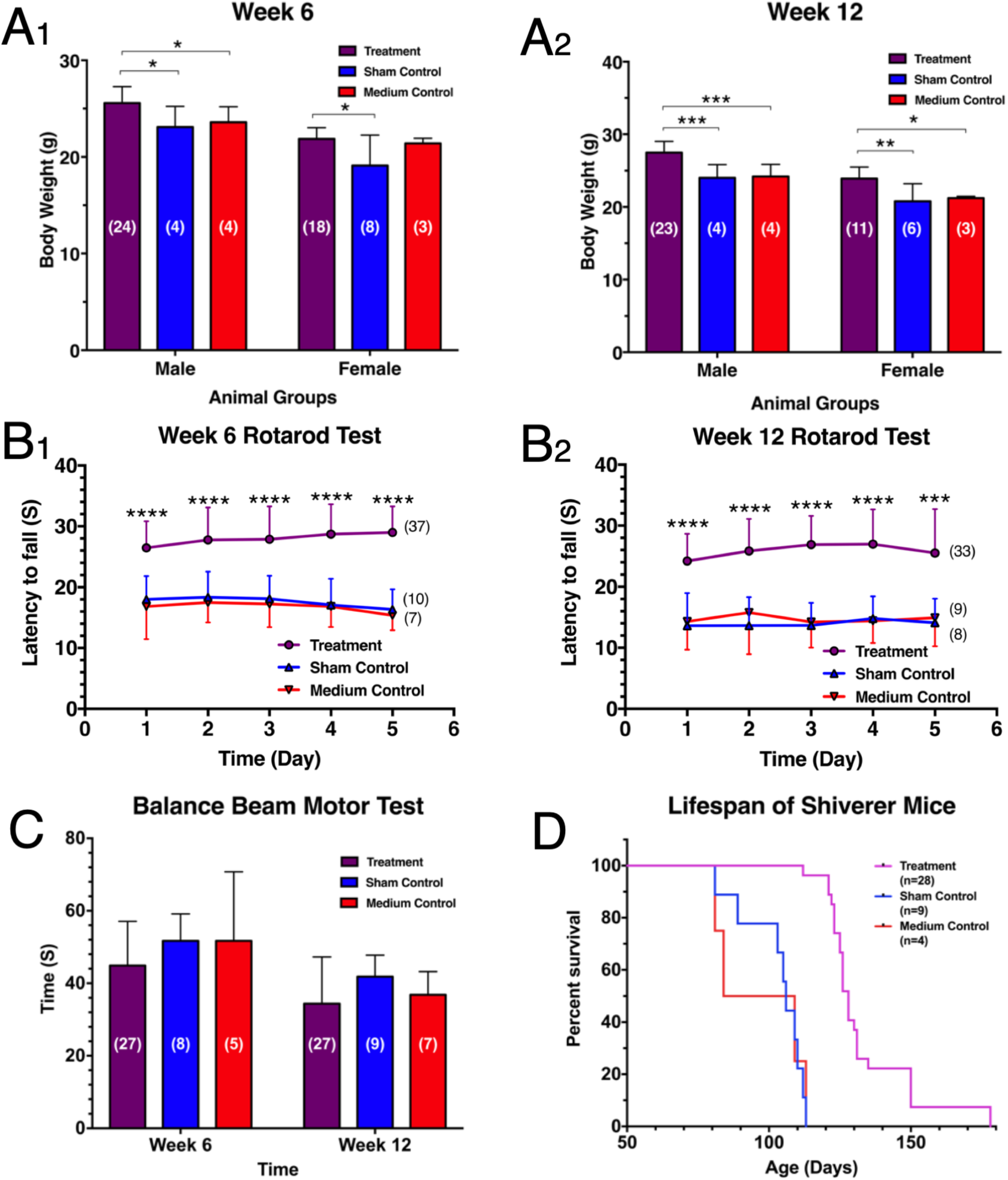
Neonatally transplanted hBM-GPCs rescue motor function, body weight and extend shiverer lifespan through remyelination. (**A**_**1**_) hBM-GPC-transplanted (treatment group) shiverer mice were significantly heavier than age-matched controls at adolescent stage (week 6, total number of mice = 61 at this time point) & (**A**_**2**_) adult stage (week 12, mean ± SD. * p<0.05, ** p<0.01, *** p<0.001 two-way ANOVA. Total number of mice = 51 at this time point). (**B**_**1**_) hBM-GPC-transplanted shiverer mice demonstrated significant better performance in accelerating rotarod motor behavioral test at week 6 & (**B**_**2**_) week 12 compared to age-matched controls. (**C**) hBM-GPC-transplanted shiverer mice performed slightly better that age-matched controls in balance beam behavioral test at week 12, but the improvement was statistically insignificant. (**D**) hBM-GPC transplantation significantly extended mean lifespan of shiverer mice to 133 days compared to 101 days of control mice.

Treated animals had significant improvements in accelerating rotarod motor behavioral test at week 6 compared to controls (Figure 5**B**_**1**_), with the mean latency of fall increased to 27.96 seconds versus 17.24 seconds in controls (a 62.84% improvement) on an accelerating profile. Treated animals demonstrated even greater improvements at week 12 (Figure 5**B**_**2**_), from 14.35 seconds to 25.88 seconds, a total of 80.34% in the motor behavioral test. In the balance beam behavioral test, treated animals were faster when traversing the beam (6.8 seconds with 13.61% increase at week 6; 5 seconds with 13.31% increase at week 12), yet due to a high standard deviation the demonstrated is insignificant (Figure 5**C**).

Treated animals also manifested significantly extended mean lifespan up to 133 days when compared with the control mean lifespan of 101 days (Figure 5**D**), giving a significant increase of 31.77%.

In conclusion, there is a significant increase/ improvement observed in: 1) body weight, 2) motor function and 3) lifespan in mice that received hBM-GPC transplantation.

## DISCUSSION

In this study, we established a novel glial induction protocol, employing the unexploited differentiation potential of human bone marrow stromal cells (hBMSCs), to rapidly and efficiently generate population of glial progenitors cells (GPCs). These bone marrow-derived glial progenitor cells (BM-GPCs) are highly enriched in oligodendrocyte progenitor cell (OPC) markers and capable of differentiating into mature oligodendrocytes in vivo.

Significant myelination was observed along anatomical regions of the motor circuitry. Extensive signals were charted at cerebellar lobules (CUL4 IV-V), the spinal vestibular nucleus (SPIV) and the external cuneate nucleus (ECU) at the hindbrain.

In midbrain sections, the corpus callosum (CC) near the injection site was most extensively myelinated. The motor cortex (MOpI, II, III) and somatosensory cortex (SSp-II1, 2/3) also demonstrated high levels of myelination. Decreasing, yet steady signals were found at the striatum (STRd) and septohippocampal nucleus (SH).

Anatomically, the corpus callosum connects the two brain hemispheres, integrating and coordinating movement between sensory and motor performances. The cerebellum regulates voluntary motor movements, including: posture, balance and coordination. The spinal vestibular nucleus regulates proprioceptive neuromuscular control of the limbs via the vestibulospinal tract, which is directed by input from the motor cortex and cerebellum. The relatively high proportions of axons myelinated reflects the myelination efficiency. Conclusively, neonatally transplanted hBM-GPCs myelinated the brain *in vivo*.

It is possible that global and dendritic synaptic input in the form of calcium signals were detected and integrated by the transplanted GPCs^46^, which hence outcompete the incapable innate OLs (of the shiverer mice), migrate and perform activity-dependent myelination along the motor circuitry. Faster and more specific signal transmission across different brain regions could translate to better behavioural output.

To bring it to a higher level, myelination of these motor function-related brain regions is likely to enhance signal transmission and hence translate into ascended motor performance. Our histological evidence is further reinforced by behavioural proof that hBM-GPCs indeed preserved function of motor circuits in mice that would otherwise suffer from significant deficits. The extended lifespan in the treatment group also hinted possible rescues from autonomic deficits through myelination.

hBMSCs have long been adopted in cell replacement strategies, being characterized as a safe and autologous source when compared to the ethically disputable embryonic stem cells (ESCs) and unstable induced pluripotent stem cells (iPSCs). We established the potential of BMSCs to be programmed into myelinating glia and shortened the induction timeframe to 14 days. This is a significant improvement from current protocols for GPC production which require at least 28 days (Ehrlich et al., 2017) ^28^. A shortened time frame enhances the translational significance of our protocol for clinical use targeting various myelin disorders.

3 patient samples of great diversity in age were all successfully derived into hBM-GPCs without modification to the protocol. This suggests that our protocol is versatile and repeatable, provided that good quality of red bone marrow (∼2-3 cm^3^) is used.

### Signaling and glial fate commitment

NG2, fully known as Neuron-Glial Antigen 2 (NG2), is a hallmark protein of oligodendrocyte progenitor cells (OPCs) (Polito et al. 2005) ^34^. NG2 is involved in cell proliferation and growth; survival, angiogenesis and cell migration. (Ampofo, et al. 2017) ^35^ It also has an interesting role to play on the fate of neuron-glial synapses during cell proliferation and differentiation.

NG2, is an integral membrane proteoglycan (Chondroitin Sulphate Proteoglycan 4/ CSPG4). We reasoned that NG2 is at the hub for controlling OL fate and maintaining OPC identity. Serving as the binding site for PDGF-AA (Tillet et al. 2002, Goretzki et al. 1999) ^36,37^, NG2 is well poised to trigger critical signaling for OPC fate commitment (Baron et al. 2000) ^38^. While its juxtamembrane domain was reported to interact with bFGF (Tillet et al. 2002, Goretzki et al. 1999) ^36,37^, which is key to keeping OPC from differentiating into mature OLs.

Yet with NG2 discovered 20 years ago, few translational studies have addressed or exploited its potential. This glial induction protocol probes the feasibility of activating PDGF and bFGF signaling through Neuron-Glial Antigen 2 (NG2). A simple growth factor-only approach was used in our protocol (detailed in Supplemental Methods), in contrast to previous studies that required transcription factor promoting viral constructs. (Ehrlich et al., 2017) ^28^ Since NG2 is also expressed in MSCs and pericytes, the success of this protocol raises the question whether all NG2^+^ cells are glial-inducible.

In previous studies (Wang et al., 2013) ^21^, OPCs derived from other cell source with different protocols were low in purity and yield (4-12%) and required follow-up fluorescence activated cell sorting (FACS) before transplantation. Most recent studies endeavored to optimize the yield (Ehrlich et al., 2017) ^28^, yet only a purity of 65-70% had been reached. Our protocol improves the purity significantly, generating a cell population with over 90% expression of OPC markers (OLIG2^+^, PDGFRα^+^, NG2^+^, SOX10^+^ and O4^+^).

Our results demonstrate the potential of using hBM-GPCs as therapy for myelin disorders. We also adopted a lower transplantation cell count (∼90,000 cells per mice) in our study and enhanced output could be expected with an increase in the quantity of hBM-GPCs engrafted. No evidence of tumorgenesis was observed in mice engrafted with hBM-GPCs up to 5 months. This suggests hBM-GPCs generated using our protocol were fate committed and safe for translation.

The study demonstrated the utility of hBMSCs as a feasible and effective source of GPCs for deriving myelinogenic oligodendrocytes. Clinical significance is further enhanced by short induction time and high purity achieved using this protocol, along with its robustness in deriving GPCs irrespective of donor age. A variety of disease settings, including congenital (e.g. myelin genetic disorders: Pelizaeus-Merzbacher Disease) and acquired myelin disorders (e.g. traumatic demyelination and multiple sclerosis) may benefit from hBM-GPC transplant.

---

**Supplementary Figure 1.**
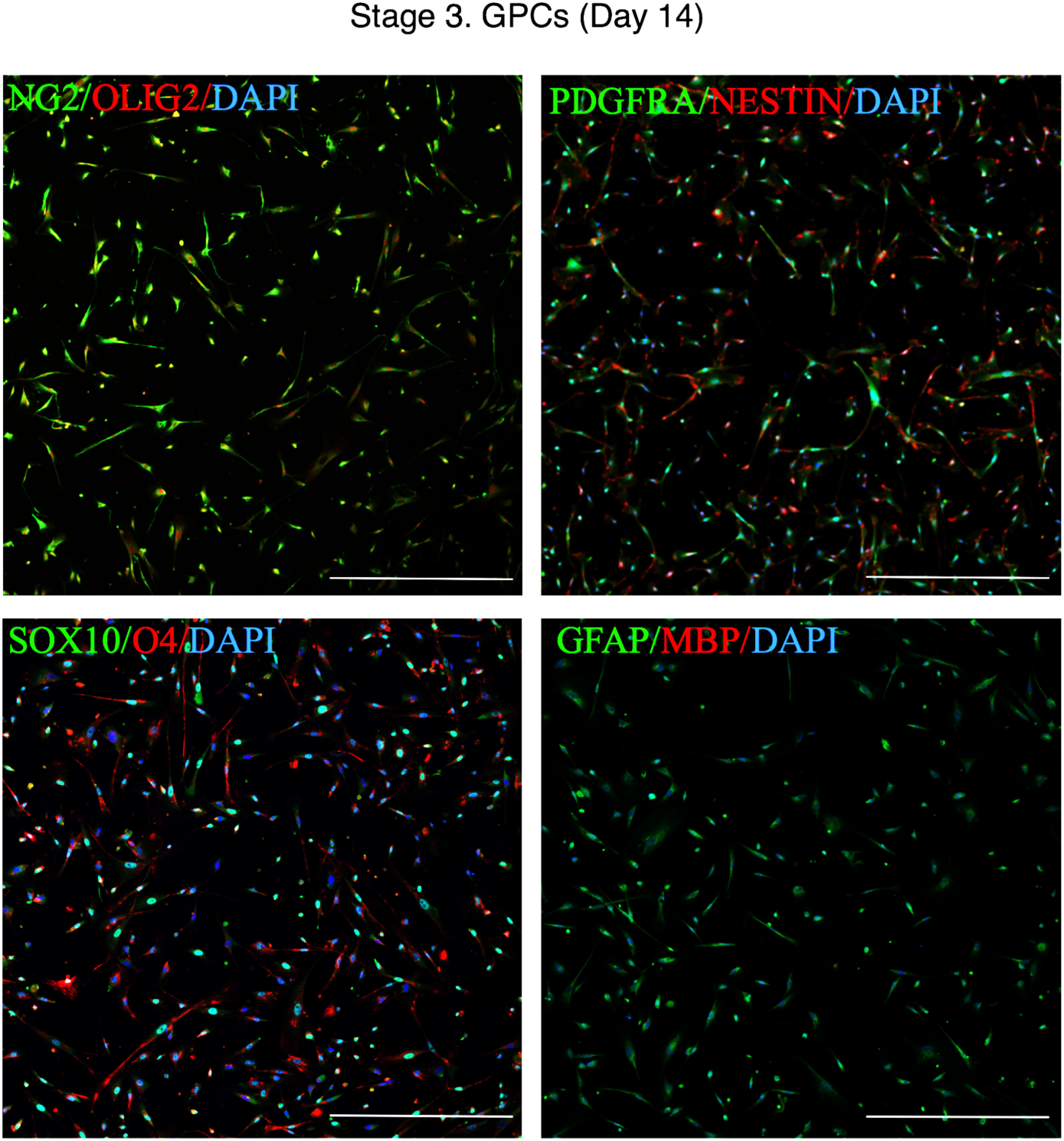
ICC Confocal Images of hBM-GPC Colonies. Low power tile scan confocal images of Stage 3 hBM-GPCs. By Stage 3, markers of early myelinogenic glial progenitors, including: NG2/ PDGFRα/ OLIG2/ SOX10/ O4 were highly expressed. Neural stem cell markers: NESTIN were also present. Mature OL marker: MBP were not expressed.

**Supplementary Figure 2.**
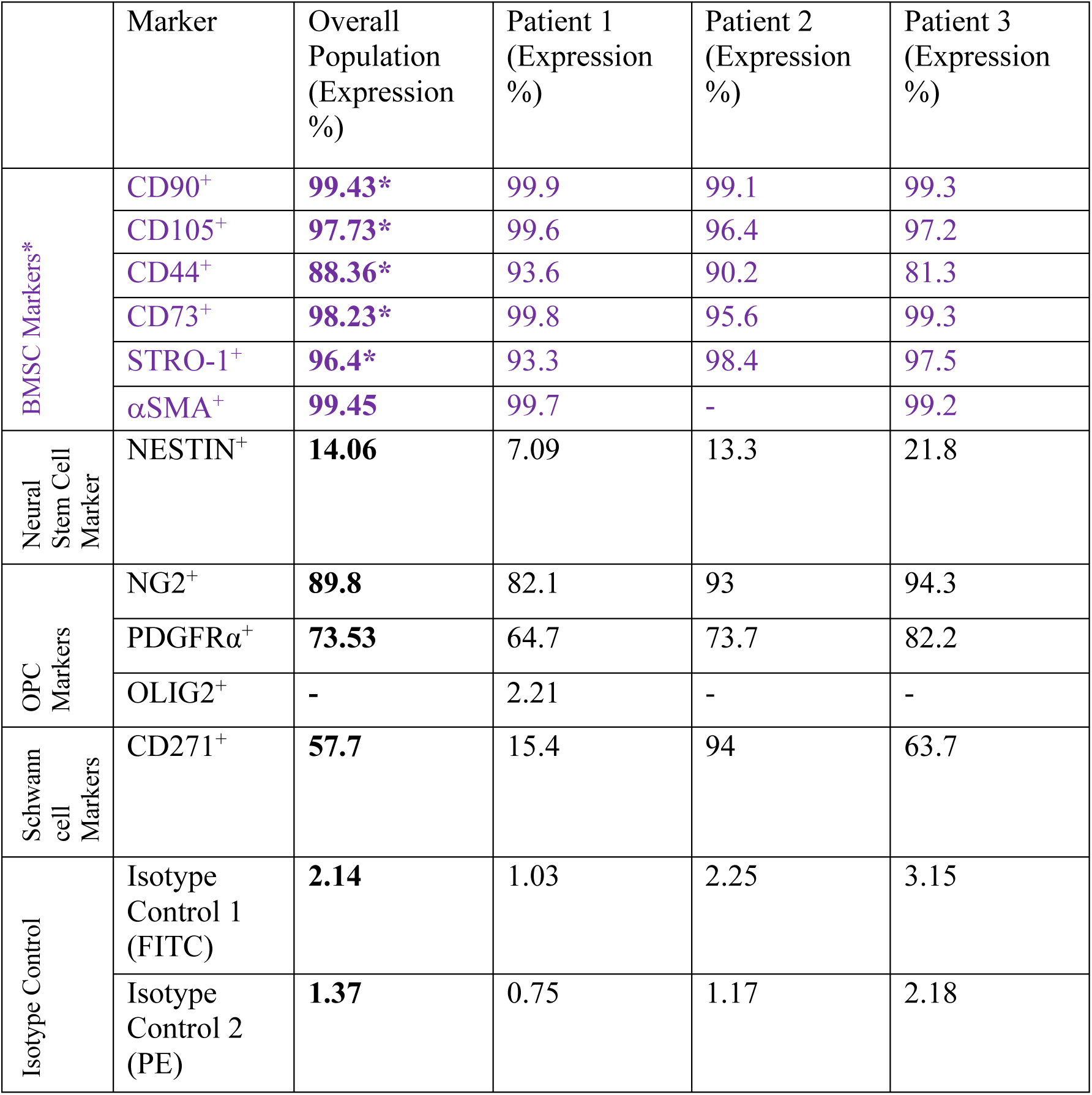
Flow Cytometric Marker Expression of Human BMSCs.

**Supplementary Figure 3.**
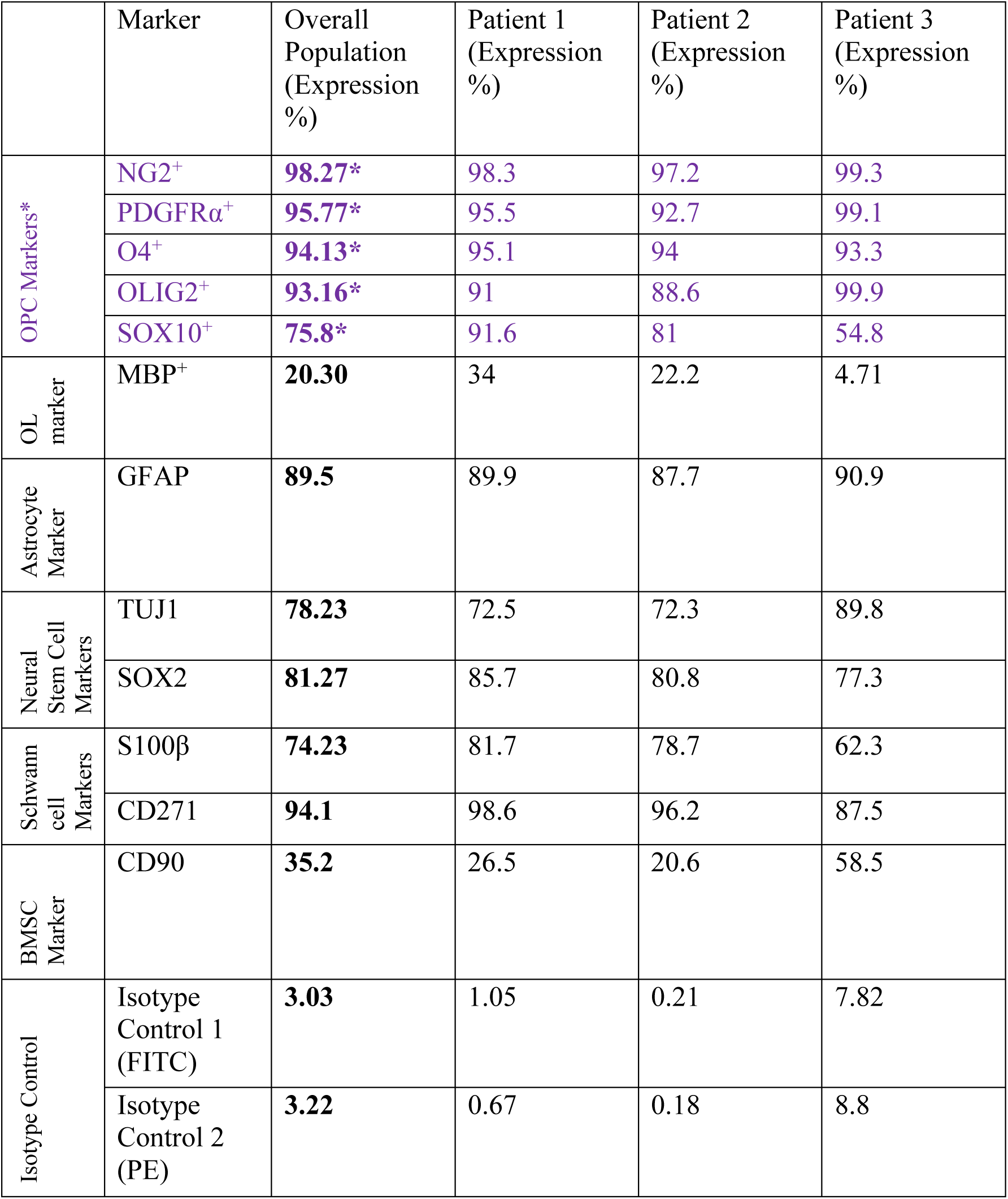
Flow Cytometric Marker Expression of Human BMSC-GPCs.

**Supplementary Figure 4.**
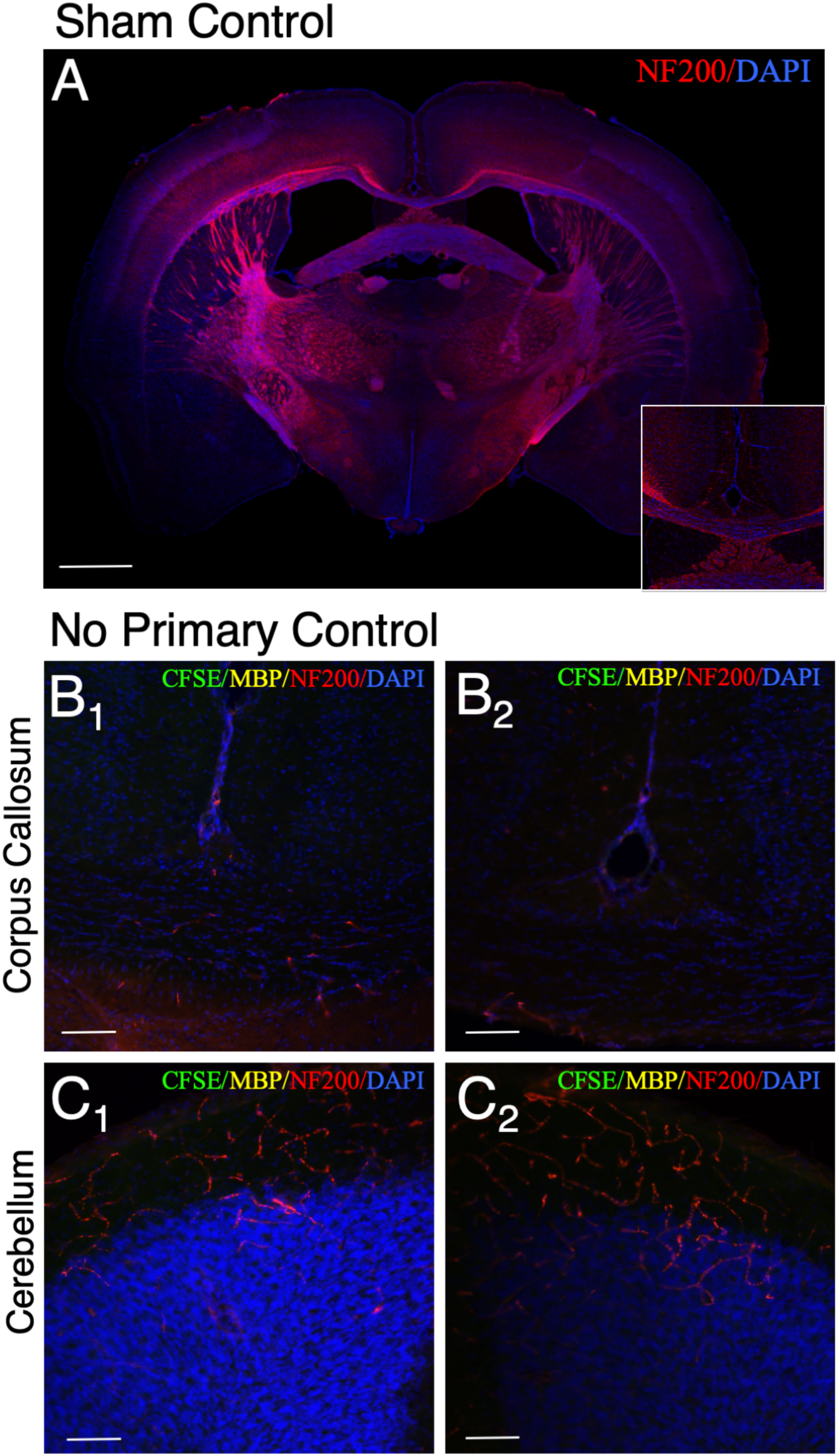
IHC Confocal Images of hBM-GPCs-derived oligodendrocytes ensheathed axons in shiverer brain (Control) Confocal images of coronal sections of the shiverer mice brain harvested at week 12. (**A**) Histology featuring the coronal mid-brain section of a sham control shiverer mouse. (Whole Brain Section) (**B**_**1**_ & **B**_**2**_) Histology featuring the corpus callosum of a treated shiverer mouse. (**C**_**1**_ & **C**_**2**_) Histology featuring the cerebellum of a treated shiverer mouse. Scale: A, B: 1 mm; C, D, E: 100 μm.

## Acknowledgements

This study was supported by the Hong Kong Health and Medical Research Fund (HMRF).

